# Cross-modal feature based attention facilitates spatial transfer of perceptual learning in motion-domain figure-ground segregation

**DOI:** 10.1101/2023.05.05.539385

**Authors:** Catherine A. Fromm, Krystel R. Huxlin, Gabriel J. Diaz

## Abstract

This study tested the role of a cross-modal feature based attention (FBA) cue on perceptual learning and spatial transfer. The trained task was figure-ground segregation in the motion domain. The experiment involved a pre-test, ten days of training, and a post-test. Twelve visually intact participants were immersed in a virtual environment and tasked with identifying the location and motion direction of a peripheral 10*^◦^*aperture of semi-coherently moving dots embedded at randomized locations within whole-field random dot motion. The aperture contained both randomly moving dots and signal dots which had global leftward or rightward motion. To manipulate motion coherence, a 3-up-1-down staircase adjusted the direction range of the signal dots in response to segregation judgments. The dot stimulus was preceded by a 1s white-noise spatialized auditory cue emitted from the fixation point (neutral group), or from an emitter moving in the direction of signal dots at 80*^◦^*/s in a horizontal arc centered on the fixation point (FBA cue group). Visual feedback indicated the selected and true aperture locations, and correctness of the motion direction judgment. Analysis measured MD discrimination within the aperture as well as segregation ability, both measured in terms of direction range threshold (DRT). At trained locations, MD DRT improved similarly in FBA and neutral groups, and learning was retained when the pre-cue was removed (ΔDRT from pretest to posttest: 61*±*10*^◦^*(SD) FBA, 74*±*10*^◦^*neutral), and transferred to untrained locations (41*±*10*^◦^*FBA, 45*±*10*^◦^*neutral). DRT for localization also improved in both groups when pre-cues were removed (49*±*10*^◦^*FBA, 44*±*10*^◦^*neutral), but only the FBA group showed full transfer of learning to untrained locations in the segregation task (32*±*10*^◦^*FBA, 23*±*10*^◦^*neutral). In summary, transfer occurred for both motion direction and segregation tasks, but the segregation transfer required the presence of the cross-modal FBA cue during training.

## 1. Introduction

The visual system is constantly faced with a barrage of information, and efficient processing of this can only come from practice and experience with interpreting the world around us. Experience shapes processing in a variety of ways, from temporary adaptation to changes in viewing conditions, like the color-change aftereffect that comes when removing tinted glasses, to the more durable changes to perception that come as the result of plasticity. Particularly in adults, plasticity occurs less often than in periods of early childhood development, but it is still possible to experience perceptual learning in the adult brain. This can come about organically, as in the case of a factory inspector becoming more accurate or quicker to spot product defects through practice, or be the result of dedicated training in a laboratory setting. The process of perceptual learning in a laboratory setting has been the subject of a great deal of research, and especially has important clinical implications in the area of visual rehabilitation (Melnick et al., 2016; Ahmad et al., 2021; Pekna et al., 2012).

Several models have been developed to understand the mechanism of perceptual learning, particularly at which stage of the visual hierarchy does plasticity occur. These models can be divided into early, mid, and late processing, are reviewed in (Watanabe and Sasaki, 2015). While there is still debate over the exact processes that govern perceptual learning, there is agreement that it is likely a complex process which can be modulated at *multiple* levels of visual processing. One of the main implications of these various models is predicting conditions which lead to either specificity or transfer of learning. Many studies where learning was shown to be retinotopically specific have lead to the theory that visual perceptual learning is modulated by early visual processing areas such as V1 (Poggio et al., 1992; Ahissar and Hochstein, 1993; Fahle and Poggio, 2002). Some models of perceptual learning explain learning as improvement to the lowest level of processing in V1 (Ahissar and Hochstein, 2004), but increasingly there is recognition that learning impacts other mechanisms on the level of read-out and combination of information at higher levels of processing (Dosher et al., 2013). Modulation at higher levels is particularly evident when training paradigms are set up to improve spatial transfer, and there are several ways to do this. The first is through the procedure known as double training, where one task is performed at a given training location and interleaved with that is a separate, irrelevant task performed in a second location. This procedure can lead to learning transfer of the primary task to the secondary location (Xiao et al., 2008; Xie and Yu, 2017; Wang et al., 2013; Talluri et al., 2015).

Another method to improve transfer of perceptual learning is through the deployment of attention. Attention can be exogenous or endogenous. Exogenous or “bottom-up” attention is generated involuntarily, often by an external stimulus, whereas endogenous or “top-down” attention is voluntarily deployed (Nguyen et al., 2020). Broadly speaking, attention can also be characterized as spatial, pertaining to a given location in a visual field, or feature-based, centered on a specific stimulus dimension such as color, direction, or orientation (Galashan and Siemann, 2017). Deployment of attention, either exogenous or endogenous, spatial or feature-based, has been shown to improve spatial transfer of visual perceptual learning (Donovan and Carrasco, 2018; Donovan et al., 2015; Hung and Carrasco, 2021). Additionally, the incorporation of cross-modal stimuli has been shown to transfer across sensory modalities (Alais et al., 2010; Shams and Seitz, 2008; Powers III et al., 2016). As perceptual learning can be modulated on different levels of visual processing, studying perceptual learning in a task which *requires* interplay between multiple levels of processing can generate new insights into the impacts of different learning paradigms on specificity. Figure-ground segregation is a mid-level visual process which requires fusion of both local and global information. It can be decomposed into several visual sub-tasks, including boundary detection, region filling and background subtraction (Huang et al., 2020) which may all depend on different mechanisms. While the primary visual cortex (V1) has shown to be involved along with area V4 in studies of non-human primates (Poort et al., 2012), the complex nature of the task requires feed-forward as well as feed-back connections to area V4 as well (Layton et al., 2014). Additionally, depending on the specific stimulus to be segregated, additional mechanisms may be involved, such as the involvement of area MT in motion-domain figure-ground segregation (Tadin et al., 2019).

Perceptual learning can improve motion-domain figure-ground segregation (Tadin et al., 2019). The plastic changes caused by perceptual learning in general can impact many levels in the visual processing hierarchy, and the exact mechanism for any given learned improvement is highly dependent on not only the stimulus used but other features of the training paradigm. In the specific case of figure-ground segregation, sound has been shown to improve mid-level visual processing and aid participants in the detection of illusory contours, a mid-level visual task which is related to figure-ground segregation (Tivadar et al., 2018). Feature-based attention has also been shown to improve figure-ground segregation (Wagatsuma et al., 2013), as well as motion direction discrimination (Martinez-Trujillo and Treue, 2004).

Our experiment measures the impact of a cross-modal feature-based attention cue on learning efficacy and learning generalizability in a figure-ground segregation task in the global motion domain, using a multi-location training paradigm. The main experimental questions addressed in this study are: 1.) Does figure-ground segregation learning occur in a multi-location training paradigm? Does motion direction discrimination? 2.) Does learning transfer to untrained locations from the figure-ground segregation and motion direction discrimination tasks? 3.) What is the effect of a crossmodal feature-based attention cue on learning, both learning rate and learning specificity? It is anticipated that the feature-based attention cue will improve learning rate as it facilitates perception in both figure-ground segregation and motion direction discrimination, as well as improve spatial transfer to untrained locations. By using a novel virtual reality test paradigm, this experiment will test whether using a figure-ground segregation task in the motion domain with a dual-task multi-location training paradigm will produce spatially general learning and improve localization ability, and also investigate whether the addition of cross-modal exogenous covert feature-based attention cue will impact learning as well as spatial generalizability.

## 2. Methods

### 2.1. Participants

Participants were recruited from the Rochester Institute of Technology (RIT) campus community. The experiment was approved by the Institutional Review Board at RIT and all participants gave informed consent prior to participation. 13 participants (4 female, 3 not reported) were recruited with normal or corrected-to-normal vision and no self-reported history of auditory processing problems. One participant self-excluded after the first session due to childhood history of amblyopia, but the remaining 12 completed the full 12 days of the experiment. The mean age in years *±* standard deviation was 21*±*2.

### 2.2. Apparatus

The experiment was conducted using the HTC Vive Pro Eye virtual reality headset. The computer used to run the experimental sessions was equipped with an NVIDIA RTX 3080 GPU and had an Intel i7-11700K CPU. The experiment was built in Unity3D, editor version 2019.4.18f1, on top of the Unity Experiment Framework (UXF) (Alais et al., 2010). Eye-tracking was done with the hardware built in to the Vive Pro Eye and controlled with the SRAnipal plugin to Unity3D, version 1.3.2.0. Auditory spatialization was done with the SteamAudio plugin, and used a generic head rotation transfer function (HRTF).

Before each experiment session, the headset was adjusted to the participant’s comfort and the headset tilt was further adjusted to ensure that the user started out viewing the display through the optical axis of the headset lenses. This was done by asking the user to change the tilt until the outer edges of the headset were sharp and clear. Once this adjustment was complete, participants followed the stock eye-tracker calibration and adjusted the headset position further if needed and adjusted the headset inter-pupillary distance (IPD) following prompts from the calibration software.

At the conclusion of the calibration sequence, a custom calibration assessment was run using a 30*^◦^* x 30*^◦^* grid of points. Participants were asked to fixate at these points in sequence and gaze position was recorded for 500ms. If average error was greater than 2*^◦^*, calibration was re-run. Average eyetracking error at the central point was 1.2*^◦^ ±* 0.6*^◦^* (standard deviation).

### 2.3. Attention cue and visual stimulus

To begin a trial, participants were asked to fixate at a central point with diameter of 0.3*^◦^* for 1000ms, and keep their head still. If the eyes or head moved more than 5*^◦^* during this period through the stimulus presentation, the trial was aborted and discarded.

After gaze was fixed on the fixation point and head was still, either an auditory cross-modal feature-based attention cue or a neutral cue to the fixation point was presented. The neutral cue was a 3D spatialized pulsed white noise cue with virtual radius of 2.5*^◦^* located at the fixation point. The pulse frequency was set to 12Hz and the cue duration was 1s. The cross-modal feature-based attention cue was the same 3D spatialized moving pulsed white noise source but rather than being stationary at the fixation point, it moved through the virtual space. Motion direction of the sound cued the forthcoming motion direction of the coherent visual stimulus within the randomly moving dot background. The auditory cue moved from left-to-right or right-to-left from one side of the participant’s head to the other. The virtual sound source moved through an arc parallel to the ground plane with radius of 0.57m so it passed through the fixation point halfway through the cue’s duration, with a speed of 80*^◦^*/s. Auditory cue parameters were set during pilot testing to produce a clear perception of motion while using a generic HRTF. Participants achieved correct auditory motion direction perception throughout the experiment.

At the conclusion of the cue period, the visual figure-ground segregation task began. The participant’s visual field was filled with randomly moving dots and within this field a 10*^◦^* diameter patch of dots moved coherently for a duration of 500ms. This coherent-motion patch was the figure to be segregated out of the ground. The randomly moving dots had a diameter of 14arcmin each, and the field had a diameter of 80*^◦^*, enough to fill the entire field-of-view of the VR headset. Dot density was 3.5 dots/*^◦^*^2^. Each dot had a lifetime of 250ms, after which it would re-spawn in a new random location. Individual dot speed was 10*^◦^*/s.

The dots within the coherent-motion patch were parameterized with two separate sources of noise. The first source of noise was the overall percentage of random dots, which was set per-individual to ensure that performance on the first day of training was similar between participants. Overall random dot noise level was set using a titration procedure (see section 2.5) on the first day of the experiment. The second source of noise was the dot direction range. The direction range of a stimulus refers to the range of trajectories the dot motion can take. For example, with dots within a motion-patch with a direction range of 0, all dots followed a path aligned with the cardinal rightward or leftward direction. Dots within a motion patch with a direction range of 80*^◦^* could take any trajectory between 40*^◦^* above the cardinal direction to 40*^◦^* below the cardinal direction. Direction range was adjusted each trial following a 3-up-1-down staircase. The combination of overall noise adjusted per-participant and direction range adjusted per-trial ensured that the task direction range reached an asymptote around the participant’s threshold for segregation.

**Figure 1:**
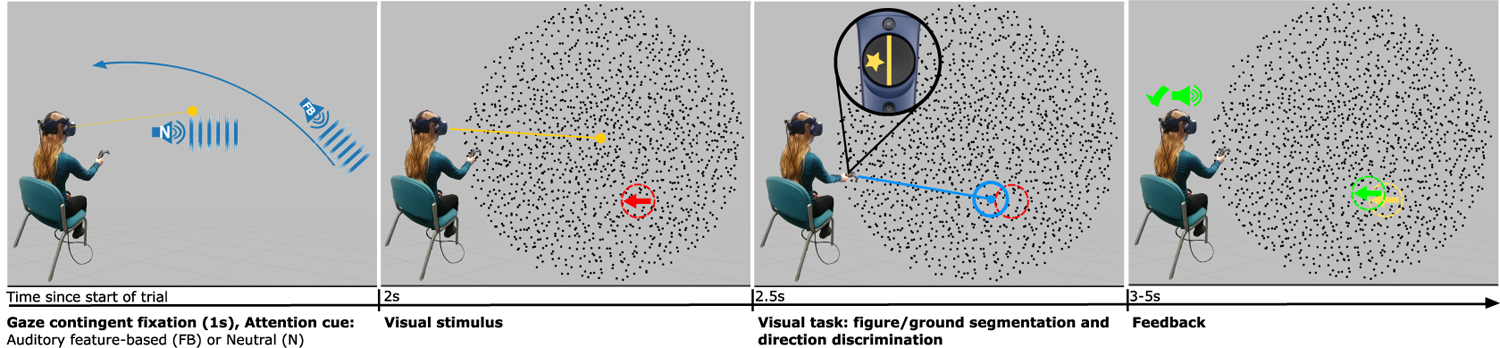
Trial Sequence In this task, a user is seated and wearing a virtual reality headset. On the start of a trial, a central fixation point is visible in a featureless grey virtual world. Each trial is gaze contingent, and begins once the integrated eye tracker detects a fixation lasting 1s. Once the fixation duration is met, the auditory attention cue plays for 0.5s. In the neutral group, the cue is a stationary pulsed white noise sound centered at the fixation point. In the feature-based attention cue group, the cue is the same pulsed white noise, but moving in an arc around the participant’s head. The direction of motion of the sound matches motion direction of the upcoming target stimulus. After the cue period ends, the field of view is filled by moving noise dots, in which a coherent target stimulus is embedded. The target stimulus is active for 0.5 seconds, whereafter the full field reverts to random motion and is present for up to 3s or until the user makes a response. The response is made by pointing to the target’s location with the handheld controller, and indicating the target’s motion direction with a button press. Feedback is then provided as a positive or negative indicator sound based on whether the localization judgment was within the cutoff range, plus a visualization of the true target location versus the indicated, as well as the true and indicated motion directions.

### 2.4. Task

After presentation of the attention cue (either neutral or cross-modal feature-based) and display of the coherent-motion stimulus, participants performed a pair of related tasks. They were instructed to indicate both the location and motion direction that the coherent motion patch possessed. This was done while the full field of random motion remained visible, to reduce the chances of a motion-aftereffect from the coherent patch. To indicate that the response period had begun, a cursor beam appeared from the handheld HTC Vive controller. This beam terminated at the point of intersection between the beam with the visual stimulus plane, and this provided participants with a visual indication of where on the plane they were pointing. Participants carried out the pair of tasks by pointing to the former coherent-motion-patch location with the controller, and indicating motion direction by pressing one side or the other of the selection pad on the controller. Pressing the direction on the selection pad also locked in the localization response to the location in which the controller was at the time of the button press.

Feedback was provided by showing the participant the true location of the coherent-motion patch versus the indicated location, as well as the true motion direction versus the indicated motion direction. A yes/no sound was also played to indicate whether localization fell within 7*^◦^* of the true location or not. 7*^◦^* was chosen as the cutoff point through pilot data showing that localizing the target within 7*^◦^* was an indication that the participant had properly segregated the figure from the ground, and any error in the localization judgment was noise contributed by other sources than figure-ground perception ability (see Section 3 for details).

### 2.5. Procedure

Each participant completed four stages of the study: random dot percentage titration, pre-test, training, and post-test. The titration and pre-test were completed on the first day, followed by ten days of training, and a post-test (identical to the pre-test) on the twelfth day as detailed in Figure 2.

**Figure 2:**
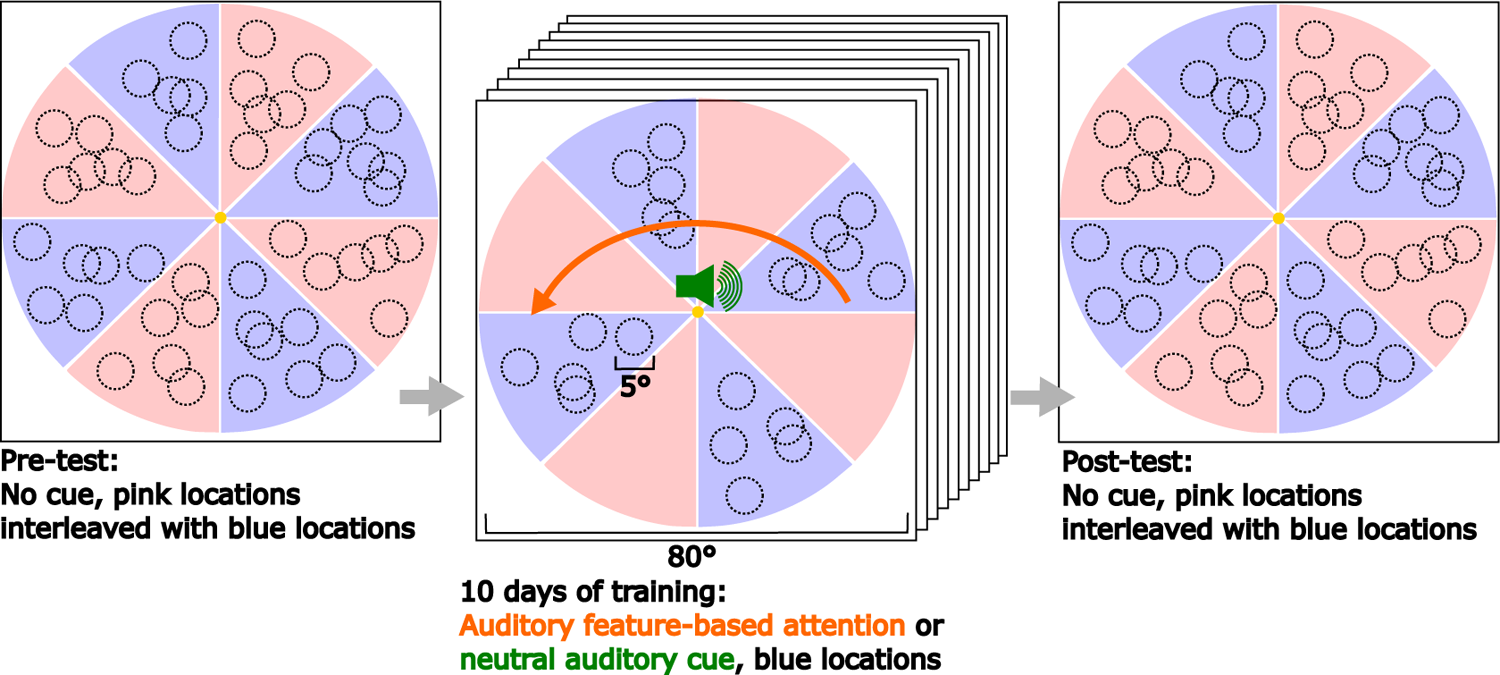
Experiment progression: The experiment consists of a ten day training period bookended by a pre-test and a post-test. During the pre- and post-tests all locations in the visual field are potential target locations, and no auditory cue is presented to either group. During the training sessions, targets may only appear in the visual field regions indicated in blue. Participants also receive an auditory pre-cue, either a neutral cue or a feature-based attention cue indicating the motion direction of the dots within the upcoming target.

In the titration block, overall noise level in the coherent-motion patch was initially set to 20% and participants were asked to complete 20 practice trials to familiarize themselves with the apparatus and tasks. If within those 20 trials, participants failed to correctly respond at least once to a trial with 80*^◦^* of direction range, random noise dot percentage was adjusted to 10% and the practice was re-started. If the participants successfully completed three trials at 120*^◦^* of direction range within the 20 practice trials, random noise dot percentage was adjusted up to 30% and the practice was re-started. This was repeated until trials at 120*^◦^* of direction range were successfully completed only once or twice during the initial 20 trials, whereupon the participant continued until 150 titration trials were completed. Thresholds were calculated on the completed 150 trials to ensure that the starting threshold was between 100 and 200 degrees of direction range for the segregation task. All participants satisfied this criteria and were able to continue the experiment.

Once the initial random noise dot level was set, participants completed a pre-test stage on the first day of the experiment. The pre-test consisted of two blocks of 300 trials each of the dual segregation/motion direction discrimination task, without any attentional pre-cues. In each block, trials were randomly presented in all locations within the visual field, including both training regions and regions of the visual field which would not be trained. Training and untrained locations were interleaved randomly to reduce the effect of practice on performance in these regions, rather than testing them sequentially. Participants removed the headset and took a break in between blocks, to avoid visual fatigue and discomfort from wearing the VR headset for long periods of time. Eye-tracking was re-calibrated each time the headset was removed and replaced. At the conclusion of training, the same test was conducted as a post-test.

The training period lasted for 10 days, in which the participants would complete 300 trials each day during a consistent time slot. During training, either the neutral cue or the cross-modal feature-based attention cue was presented ahead of each trial. The coherent-motion patches also only appeared in select “training locations” (see figure 2). When interviewed during the experiment debrief, participants reported no conscious knowledge that training was conducted in a restricted set of locations, or that it differed from the pre- and post-test in any way save for the presence of the pre-cue.

### 2.6. Analysis

Broadly, the analysis focuses on comparing the performance in the pre-test and post-test as well as assessing the impact of training within the training period. With the dual segregation/motion direction discrimination task, several axes of performance were examined to address the experimental questions posed by this study.

Firstly, if global motion direction judgment improves, we will see this as an increase in the direction range threshold for motion direction (DRTmd). This threshold is calculated from a psychometric function representing the percent of correct direction judgments at each level of direction range. Thresholds are calculated with a 5% lapse rate and a 50% chance level. This means that even with perfect perception, 5% of the trials will be random mistakes, and even with a random guess, 50% of the trials will still be guessed correctly, as the choice is only between two options see Figure 3 panel D.

**Figure 3:**
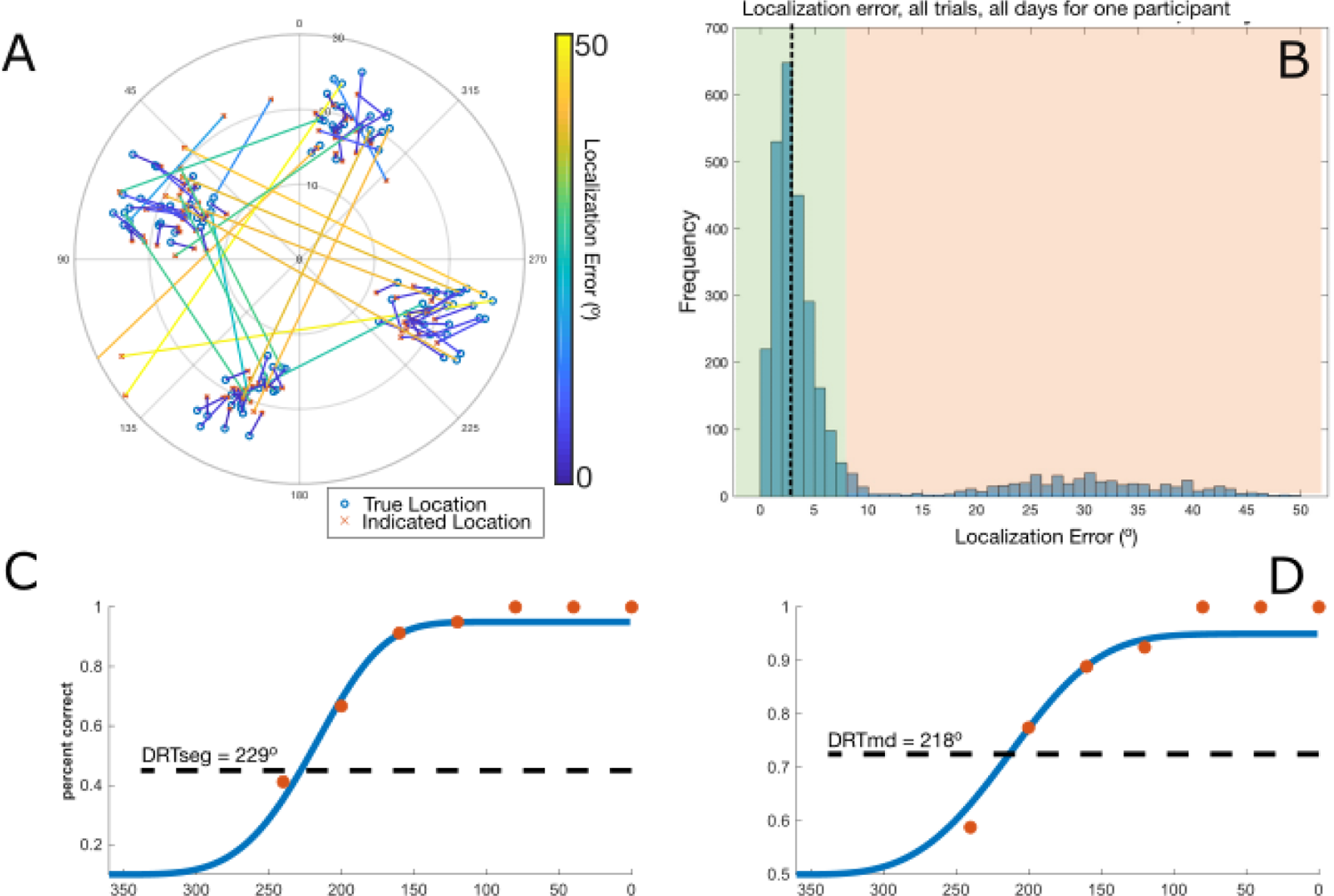
Analysis procedures A: Representative localization judgments from a selection of training trials. True location shown as blue circles, and the participant’s indicated location shown as red crosses. Error vectors connecting true and indicated locations are color coded based on error size, with 50*^◦^* of visual angle shown in yellow and 0*^◦^* shown in violet. B: Frequency plot of all errors for one participant, over all days. This distribution is strongly bimodal for all participants, and the cutoff for a “correct” judgment is shown highlighted in green. C: Sample Weibull function fit of DRTseg for localization responses, with thresholds determined at 52.5% correct, halfway between 95% correct and the chance level of 10% correct. D: Sample Weibull function fit of DRTmd for motion direction judgments, with thresholds determined at 72.5% correct, halfway between 95% correct and the chance level of 50% correct.

Second, if segregation ability improves, we will see this as an increase in the direction range threshold for segregation (DRTseg). This threshold is calculated similarly to the DRTmd, from a psychometric function representing the percentage of correct segregation judgments at each level of direction range. However, the response to the segregation task is collected as a continuous measure of localization error, and must be transformed to a binary correct/incorrect judgment before percentage correct can be calculated. Though the measure is continuous, the underlying responses show a strong bi-modal distribution with a clear cutoff point between trials where the target was perceived and when the participant was making a random guess. This cutoff point was reinforced by feedback during the experiment which was positive when segregation judgments fell within the cutoff region, and negative when they were outside it (see Figure 3 panel B). Once the psychometric function was created with the percentage of trials within the cutoff point on the vertical axis and direction range of those trials on the horizontal axis, thresholds were calculated using a 5% lapse rate and a 10% chance level (see Figure 3 panel C). As in the DRTmd measure, there was a 5% chance that even with perfect perception the participants would make a random error, and with the geometry of the task (size of field of view, size of target stimulus and response cursor) there was a 10% chance that any random guess as to the target location would fall within the cutoff point and be counted as correct.

Third, if localization accuracy and precision improve with training, this will be observed as a reduction in average localization error as well as a reduction in the variance of localization error. Localization error refers to the distance between the centers of the target and the response cursor in units of degrees of visual angle. To avoid confounding this measure with random guesses, (in which participants are trying to localize a target which they incorrectly perceived and does not exist) only trials which fall within the DRTseg cutoff point are analyzed. This means that they were properly segregated from the background. Additionally due to the staircase design of the experiment, most trials are conducted near the DRTseg threshold. To make a fair comparison of localization judgments, segregation ability should be about the same for each localization judgment being compared. Thus, only trials from direction ranges near the DRTseg on a given day will be compared. To sum up: localization error is filtered using the same cutoff point as in the DRTseg calculation, then further filtered to only include trials at direction ranges around the calculated DRTseg. These filtered trials are then averaged to get localization accuracy, and the standard deviation is calculated to get localization precision.

With these three analysis measures, learning transfer to trained and un-trained locations is also tested. If improvement on any measure (DRTmd, DRTseg, localization accuracy and localization precision) transfers to the un-trained locations, the change from pre-test to post-test should be the same in the training locations as it is in the untrained locations.

Similarly, the effect of the cue is also tested. If the addition of the cross-modal feature-based attention cue improves performance on any measure (DRTmd, DRTseg, localization accuracy and precision), the change from pre-test to post-test should be greater in the group trained with the cross-modal feature-based attention cue versus the group trained with a neutral cue. If the cross-modal feature-based attention cue leads to more generalizable improvements, then the difference in change from pre-test to post-test in the trained versus untrained location should be smaller for the group trained with the cross-modal feature-based attention cue than for the group trained with the neutral cue.

#### 2.6.1. Statistical tests

Tests related to generalizability and group level effects were conducted using a linear mixed effects model within the R statistical software environment (version 4.0.5 (R Core Team, 2021)). DRT thresholds were the response variable. Fixed factors were test day (pre-test or post-test), presence of cue during training, testing locations (trained and untrained), and type of threshold (DRTseg or DRTmd). The participant was fit as a random factor. This random factor contributed 45% of the overall variance in the model. This model was then used to calculate estimated marginal means, from which interaction contrasts within the training day were computed in post-hoc testing to determine possible differences in the change from pre-test to post-test over combinations of these factors.

## 3. Results

This study produced learning in both groups, which in the case of the audiovisual group fully transferred to the task condition with no attention cue. Spatial transfer of learning was observed for both groups in the DRTmd measure, but learning in the DRTseg measure only transferred to untrained locations when training was done with the auditory feature-based attention cue.

During the training period, both groups showed significant learning in the DRTseg measure, as shown in Figure 4. The auditory feature-based attention cue group performed near ceiling in the DRTmd measure for the duration of the training period, as the auditory pre-cue gave full knowledge of the motion direction at every trial. In the neutral cue group, there was also improvement in the DRTmd measure. The learning observed in the DRTmd measure was not significantly different from the learning in the DRTseg measure.

**Figure 4:**
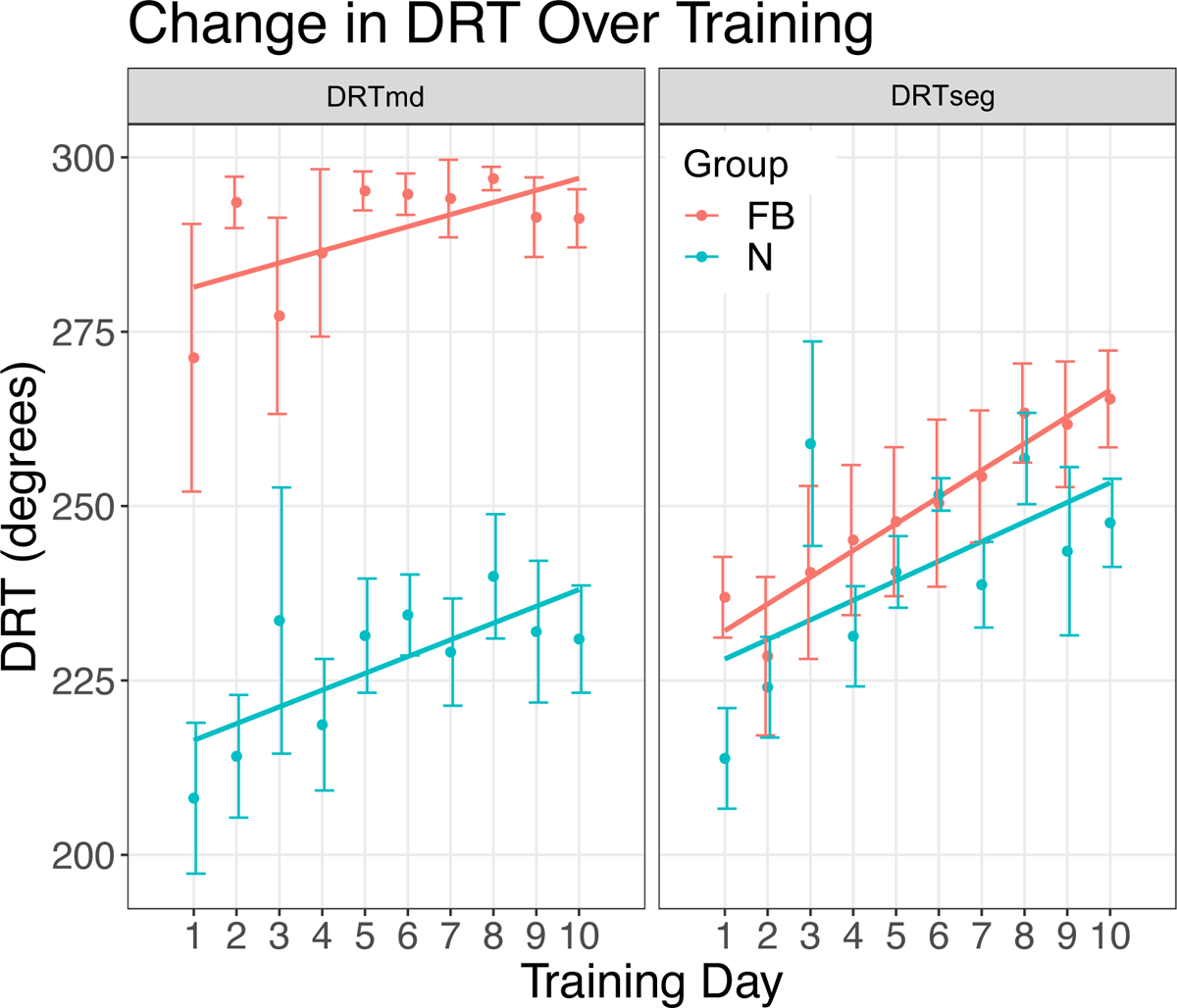
Training results. Points shown represent each DRT measurement averaged over all participants within the group, error bars are standard deviation. The horizontal axis is day of training. In the left panel, the vertical axis is DRTmd, and in the right panel and the vertical axis is DRTseg. The DRTmd measure shown in the left panel was consistently near ceiling for the auditory feature-based attention cue group (orange). The neutral cue group (blue) had a slope estimated at 2.43*^◦^/*day with standard error 1.05*^◦^/*day. In the DRTseg measure shown in the right panel, the feature-based attention group (blue) showed a slope estimated at 3.8*^◦^/*day with standard error 0.83 and the neutral group had a slope estimated at 2.9*^◦^/*day with standard error 0.84. These two slopes were not significantly different from one another. Additionally, the slopes over DRTmd and DRTseg within the neutral groups were not significantly different from one another

**Figure 5:**
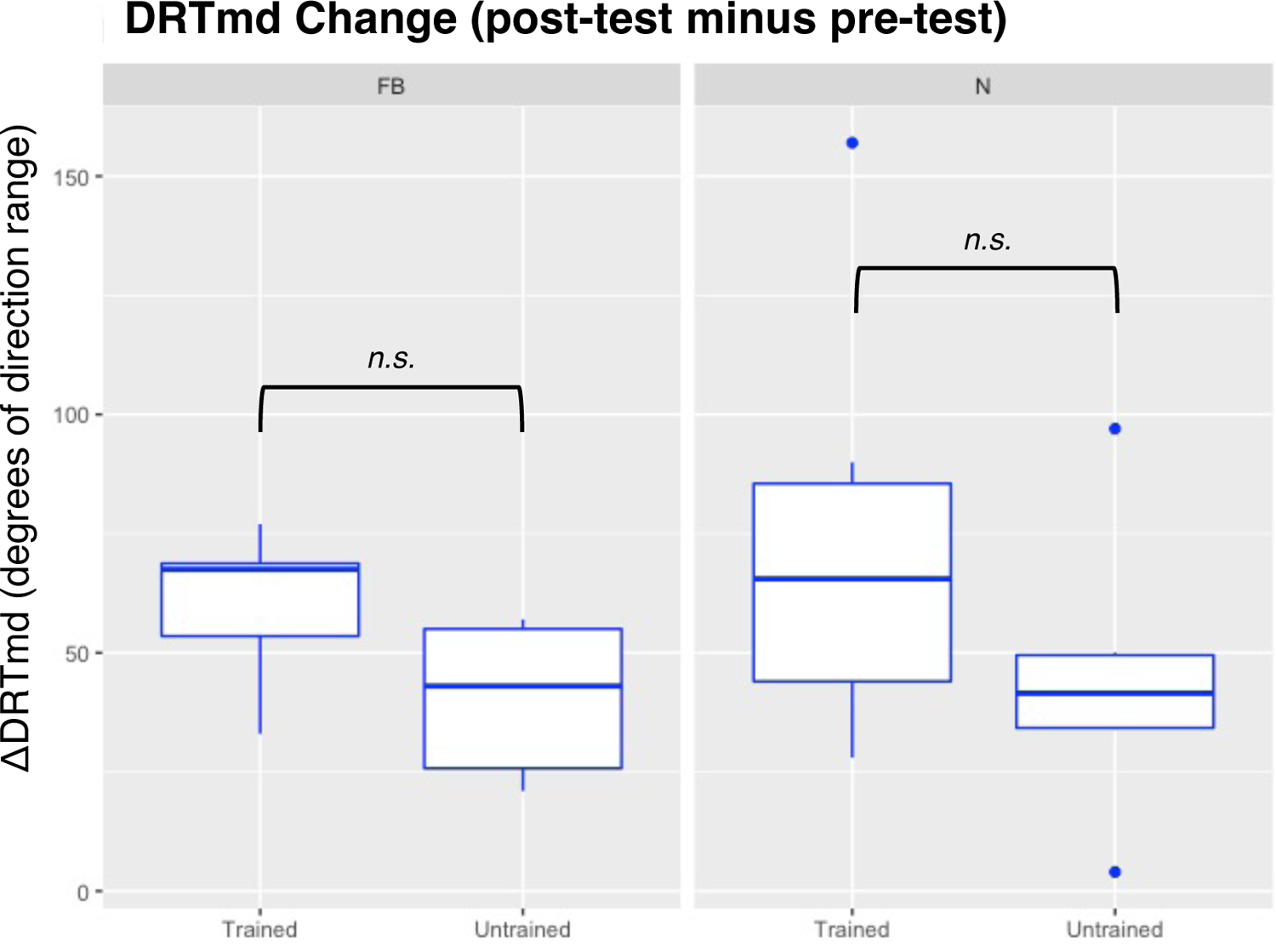
Pre- to post-test change in motion discrimination judgments (DRTmd) The feature-based cue group (left panel) improved an estimated 60.5*±*10.4 degrees of direction range in the trained locations (left box plot), and 40.5*±*10.4 degrees of direction range in the untrained locations (right box plot). These values were not significantly different from each other, indicating full transfer of DRTmd learning in the feature-based cue group. The neutral cue group (right panel) improved an estimated 74.2*±*10.4 degrees of direction range in the trained locations (left box plot), and 44.7*±*10.4 degrees of direction range in the untrained locations (right box plot). These values were not significantly different from each other, indicating full transfer of DRTmd learning in the neutral cue group.

**Figure 6:**
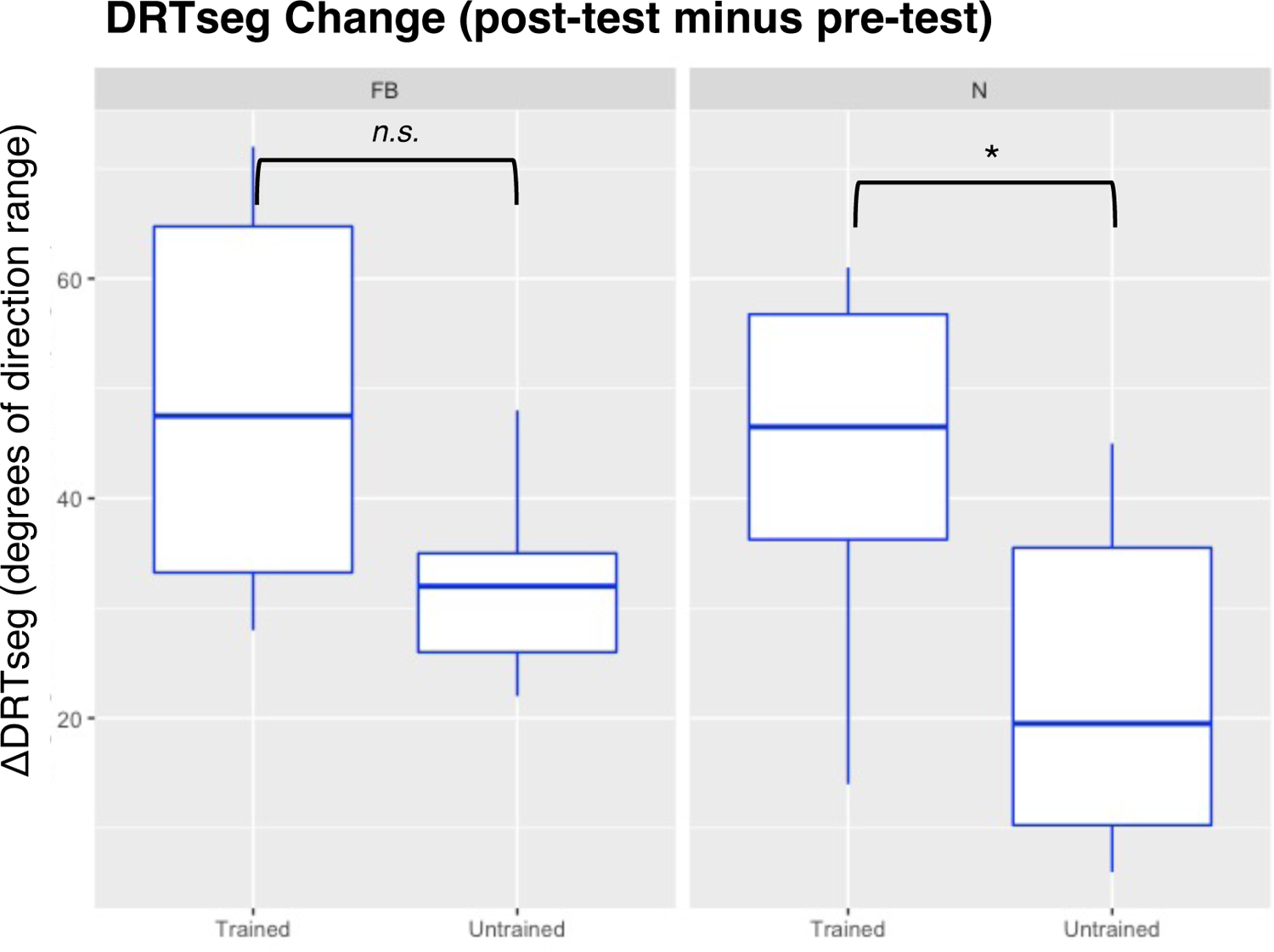
Pre- to post-test change in figure-ground segregation threshold (DRT-seg) The feature-based cue group (left panel) improved an estimated 49.0*±*10.4 degrees of direction range in the trained locations (left box plot), and 32.3*±*10.4 degrees of direction range in the untrained locations (right box plot). These values were not significantly different from each other, indicating full transfer of DRTseg learning in the feature-based cue group. The neutral cue group (right panel) improved an estimated 43.5*±*10.4 degrees of direction range in the trained locations (left box plot), and 23.0*±*10.4 degrees of direction range in the untrained locations (right box plot). These values were significantly different from each other (*p* = 0.004), indicating partial transfer of DRTseg learning in the neutral cue group.

The main effect of the pre-test to post-test was highly significant (*p <* 0.001) in the linear mixed effects model. No interactions between day and any other effects were significant. Within the main effect of pre- to post-test change, contrasts were calculated to determine the magnitude of change for each group, in each set of locations and for each measure.

To evaluate transfer of learning to the task without the presence of the auditory feature-based attention cue, change in performance from pre-test to post-test in the training locations on both the DRTseg and DRTmd measures was calculated and compared between groups. In the training location, the feature-based cue group had a significant (*p <* 0.001) estimated increase in the DRTmd measure of 60.5 degrees of direction range with a standard error of 10.4 degrees of direction range. The neutral cue group showed a significant increase (*p <* 0.001) in the DRTmd measure with estimated improvement of 74.2 degrees of direction range and standard error of 10.4 degrees of direction range. In the audiovisual group, we observed a significant (*p <* 0.001) increase in the DRtseg measure estimated at 49.0 degrees of direction range with a standard error of 10.4 degrees of direction range. For the neutral cue group, the DRTseg also increased significantly (*p <* 0.001) with an estimated improvement of 43.5 degrees of direction range and standard error of 10.4 degrees of direction range.

In the untrained locations, the feature-based cue group saw a significant (*p <* 0.001) improvement in the DRTmd estimated at 40.5 degrees of direction range with standard error of 10.4 degrees of direction range, and significant (*p* = 0.003) improvement in DRTseg estimated at 32.3 degrees of direction range with standard error of 10.4 degrees of direction range. The neutral cue group had significant (*p* = 0.001) improvement of DRTmd estimated at 44.7 degrees of direction range, standard error of 10.4, and significant improvement of DRTseg estimated at 23.0 degrees of direction range with standard error of 10.4 degrees of direction range.

Post-hoc tests were then done to estimate the differences between contrasts. One notable result from this study is that motion-direction discrimination learning, measured with DRTmd, transfers to untrained locations, regardless of the presence of the attentional cue. The contrasts comparing pre-test to post-test change between trained and untrained locations for the DRTmd measure show there is no significant difference in any of the changes from pretest to post test, for the trained and untrained locations. This indicates that both groups displayed full transfer of motion direction discrimination learning.

However, we do see that the presence of the cue during training impacts the transfer of *figure-ground segregation learning*. In the group which received the feature-based cross modal attention cue during training, learning in the DRTseg measure transferred fully to the untrained locations. There was no significant difference in the improvement in performance for the trained locations versus the untrained locations. This differed in the neutral cue group. While there was significant learning in both the training and untrained locations, the comparison between these changes showed a significant difference (*p* = 0.004) between the learning in the training and untrained locations. The neutral cue group still showed transfer of learning to the untrained location, but it was a smaller improvement than observed in the training location, indicating partial transfer.

The other results from this study are that the perceptual learning measured by DRTseg and DRTmd do not impact localization accuracy and precision as shown in Figure 7 and Figure 8. Localization accuracy and precision were each fit with a separate linear mixed effects model including separate models for the change over the training period and the change between the pre-test and the post-test. In all models, the response variable was either accuracy or precision, and the random effect was the participant’s study ID. The fixed effects were day of the study and group assignment for the model of the training days, and the models for the pre- to post-test data also included the locations as a fixed effect. These models showed no significant effects over any of the fixed factors or interactions between them.

**Figure 7:**
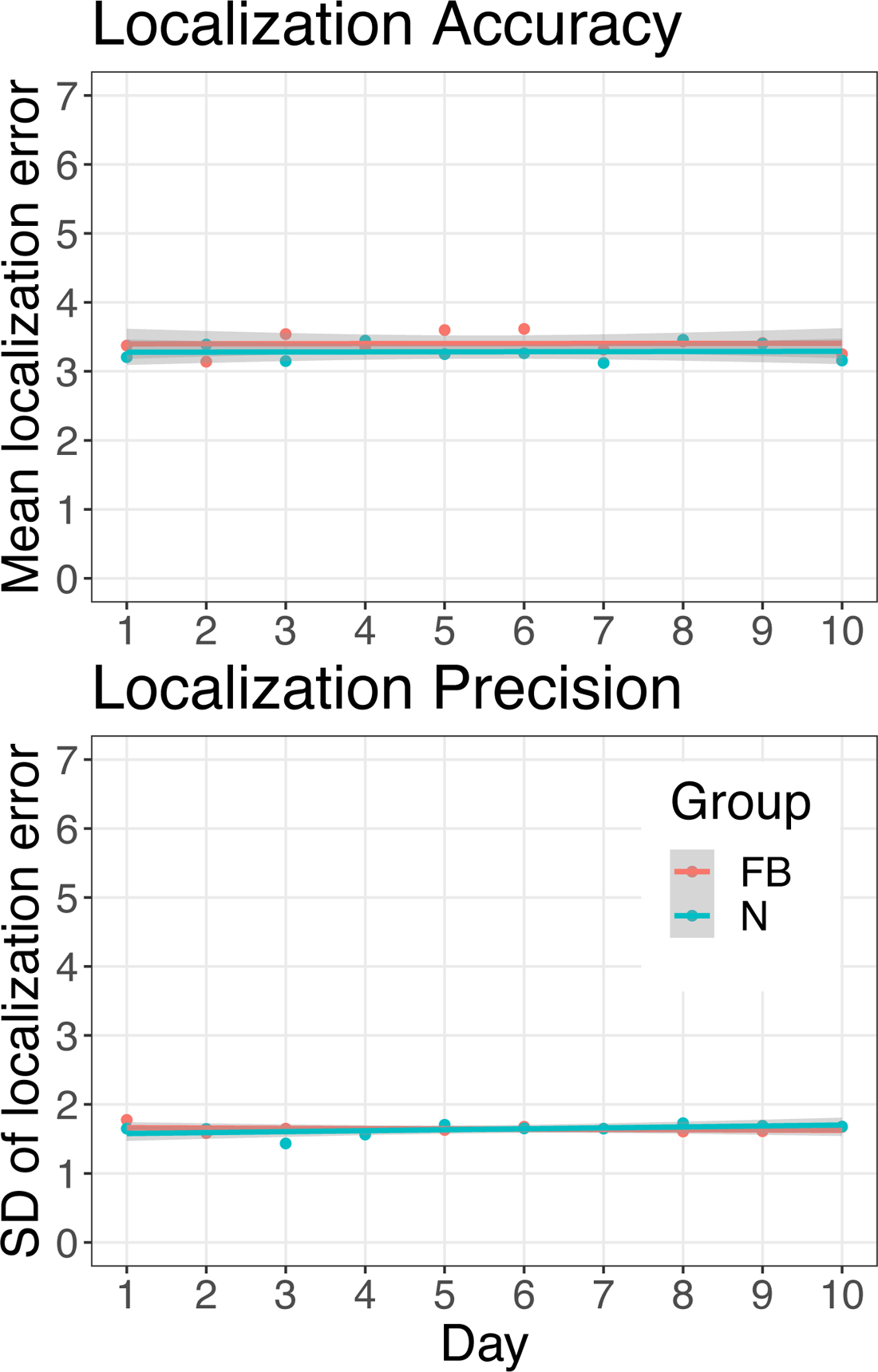
Training localization accuracy and precision. The top panel shows the localization accuracy, or average localization error, averaged over all participant on each of the ten days of training. The bottom panel shows the localization precision, or standard deviation of localization errors, averaged ov1e8r all participant on each of the ten days of training. For both groups, neither accuracy or precision had significant change over the training period.

**Figure 8:**
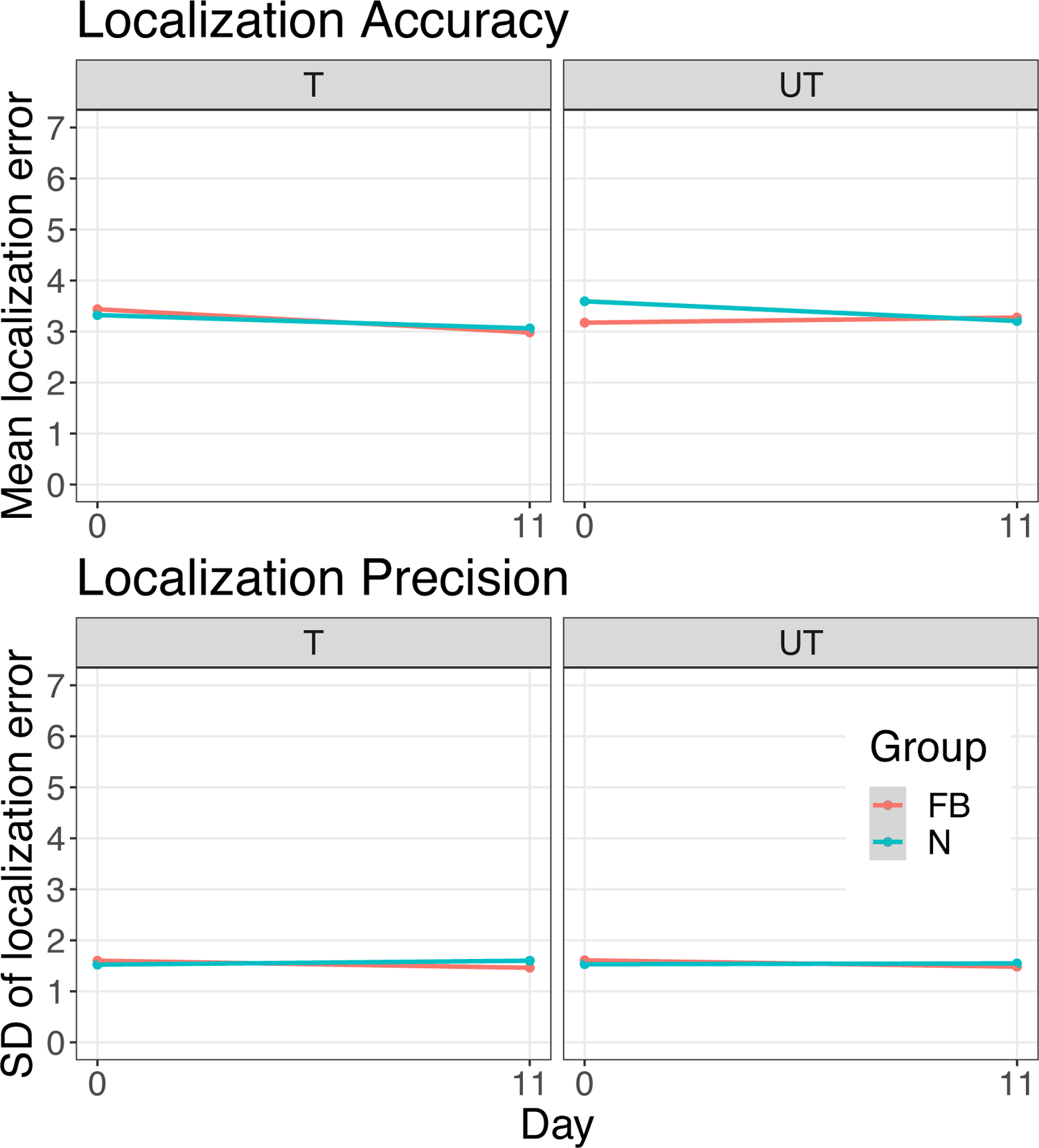
Pre- to post-test localization accuracy and precision, in training and untrained locations. The top panels show the localization accuracy, or average localization error, averaged over all participant on each of the ten days of training. The top right panel is trials in the training locations, and top left panel is untrained locations. The bottom panels show the localization precision, or standard deviation of localization errors, averaged over all participant on each of the ten days of training. The bottom right panel is trials in the training locations, and bottom left panel is untrained locations. For both groups and in both sets of locations, neither accuracy or precision had significant change from pre-test to post-test.

## 4. Discussion

This task produced tangible learning results in the dual-task paradigm of figure-ground segregation and motion direction discrimination learning. The presence of the auditory, feature-based attention cue mainly impacted transfer to untrained locations. The motion direction discrimination learning transferred fully in both groups, regardless of the presence of a feature-based attention cue. However, the figure-ground segregation learning only fully transferred to untrained locations with the addition of the feature-based attention cue. These results speak to the multiple levels of processing on which perceptual learning can operate.

First of all, in the trained locations, both groups showed significant learning on both the figure-ground segregation task as well as the motion direction discrimination task. This learning occurred strongly even though the training took place in a spatially-jittered manner, where the location of the training stimuli varied constantly during the session. While training was only performed in half the visual field, split up into 4 wedges out of 8 (see Figure 2), participants reported no conscious knowledge that locations were constrained during training. This result is of particular interest in the context of rehabilitation, as it shows that fewer trials in any given location can still produce strong learning. This is in alignment with other studies of double training (Xiao et al., 2008; Mastropasqua et al., 2015) in which stimulation of multiple retinal locations during training can lead to full spatial transfer at locations with comparably few trials. Though our method of randomized stimulus placement differs from the conventional double training procedure where only two locations are trained, the same principles central to double training apply where the attention is distributed to multiple locations and learning can accumulate across all trained locations.

This learning transferred to the visual-only pre- and post-tests for both groups despite the fact that the testing sessions were carried out with no pre-cue. Both groups demonstrated full spatial transfer of the motion direction discrimination task to the untrained locations. This could be due to several factors. Firstly, the same “double training” effects which allowed learning to occur with fewer trials in any one location may have carried over to fully untrained locations. Because participants did not consciously know the training locations only occurred in half the visual field, higher level processes such as the distribution of spatial attention may have been deployed which are known to increase spatial generalizability (Donovan and Carrasco, 2018). Crucially, however, the same transfer did not occur for the figure-ground segregation task in the neutral cue group. This decoupling of two stages has been noted in other figure-ground segregation tasks. (Yi et al., 2006; Wagatsuma et al., 2018). If training in multiple locations was solely responsible for the spatial transfer, one would expect to see the effects of this apply in both tasks, as the training method was identical for the two tasks. More likely, the learning of the motion direction discrimination task is mediated by changes in the higher levels processing needed to complete the task, and can transfer to untrained locations (Zhang and Li, 2010; Zhang et al., 2019). These higher levels in the visual hierarchy are not spatially specific (Dosher et al., 2013) and can modulate signals from multiple locations. Important to note is that visual perceptual learning has been increasingly viewed as a whole-brain process (Maniglia and Seitz, 2018) rather than mediated by a single specific mechanism. In this task, the motion direction discrimination is linked to the figure-ground segregation in that improvement on motion direction discrimination ability *could* improve overall figure-ground segregation ability, but is not a requirement. This is because figure-ground segregation relies on input from several sub-tasks. In a task like figure-ground segregation where performance is dependent on the successful completion of sub-tasks at multiple levels of pooling, perceptual learning in this task could be achieved through modulation at any of these stages. Similarly to how a chef can improve her cooking in a number of ways, at the lowest level by using fresher ingredients, next by choosing tastier flavor combinations, and finally by using better cooking techniques, any of the changes alone can result in better testing food even in the absence of the others. So perceptual learning in the domain of a complex tasks like figure-ground segregation can be accomplished through improvement on any individual sub-tasks. In our neutral group, the mechanism for improving figure-ground segregation was likely independent of the mechanism which improved motion direction discrimination, as the motion direction learning transferred to the untrained locations where the figure-ground segregation learning did not. This is supported by other studies indicating that background subtraction happens in early vision even at the level of the retina (Ölveczky et al., 2003).

Another key result from this study is that the addition of the auditory cross-modal feature-based attention cue lead to full spatial transfer of the figure-ground segregation task. This is consistent with other results showing that visual endogenous feature-based attention improves spatial generalizability of perceptual learning (Hung and Carrasco, 2021). Specifically in figure-ground segregation, feature-based attention has been shown to improve early visual processing in figure-ground segregation (Wagatsuma et al., 2013).

Our study shows that modulation of auditory feature-based attention has implications for the transfer effects of visual perceptual learning. This result is also interesting in the context of visual rehabilitation. In cortically blind participants, feature-based attention has been shown to generate recovery of fine direction discrimination abilities (Cavanaugh et al., 2017). The use of a cross-modal attention cue could be particularly beneficial for this patient population as other studies have shown promising rehabilitation through the use of other cross-modal stimuli (Purpura et al., 2017; Cioni et al., 2015; Cappagli et al., 2017). Presenting this feature-based attention cue in the auditory modality also raises some interesting questions about the mechanisms involved in cross-modal attention and perceptual learning. Area MT has been shown to be heavily involved in visual motion processing (Yates et al., 2020), as well as in figure-ground segregation (Tadin et al., 2019). Additionally, though area MT is not typically thought of as being involved in auditory processing, this area has been shown to be capable of responding to auditory motion stimuli in patients with early blindness (Saenz et al., 2008) as well as in early blind adults who have learned to echolocate (Thaler et al., 2014), indicating that cross-modal plasticity may be possible in this area. Even in studies of visually intact participants, this region of the brain has been shown to represent auditory motion direction along with visual motion direction (Rezk et al., 2020). This may be a possible mechanism for the cross-modal transfer seen in the improved performance even in the absence of the auditory cue during the visual-only post-test. While the receptive field size in area MT is larger than in V1 (Amano et al., 2009; Felleman and Kaas, 1984), it is not large enough to fully explain the spatial transfer of learning to untrained locations especially transfer to the opposite hemifield. However, since this task paradigm used multiple training locations spatially jittered throughout the visual field, this spatial transfer may instead be impacted by the use of multiple training locations in close proximity to the untrained locations.

In conclusion, this study has shown that motion direction discrimination can be improved by perceptual training in multiple locations, and that this learning can fully transfer to untrained locations. Similarly, the figure-ground segregation ability which may use motion direction discrimination as a precursor to figure-ground segregation in this task can also be improved by perceptual training in multiple locations, but learning does not fully transfer to untrained locations. However, the addition of an auditory cross-modal feature-based attention cue can cause full spatial transfer of learning for the global motion direction discrimination task. Future work is necessary to determine whether this effect is restricted exclusively to cross-modal cues or if the transfer effect is preserved with an intra-modal visual feature-based attention cue. Additionally, further study is needed to quantify the impact of training at multiple locations. Virtual reality uses an expanded field of view and the addition of integrated eye tracking makes it easier to provide stimulus in multiple retinal locations while controlling for eye movements.

